# KneEZ Clear, an Effective Tissue Clearing Protocol to Study Musculoskeletal Tissues in the Mouse

**DOI:** 10.1101/2025.08.13.670136

**Authors:** Julia Younis, Taeyong Ahn, Luis Tovias, Azeez O. Ishola, Carolina Leynes, Jason M. Kirk, Sienna K. Perry, Akash Gandhi, the RE-JOIN Consortium, Joshua J. Emrick, Brendan H. Lee, Joshua D. Wythe, Nele A. Haelterman

## Abstract

Wholemount, 3-dimensional (3D) tissue imaging holds significant promise for analyzing heterogeneous musculoskeletal tissues, such as knee joints, that demand time- and labor-intensive processing using traditional histological methods. Current musculoskeletal clearing protocols rely on either solvent-based tissue clearing, which substantially alters the size and architecture of cleared tissues, possibly compromising downstream quantification and perhaps more importantly reducing signal from endogenous fluorescent reporters, or on expensive and time-consuming hydrogel-based approaches that requires specialized equipment. While aqueous-based clearing overcomes these challenges, there is a clear need for a method that is optimized for clearing musculoskeletal tissues and that can easily be implemented in a standard lab environment. Here, we present KneEZ Clear, a simple, rapid, and flexible aqueous-based method that renders mineralized and non-mineralized tissues of murine knee joints optically transparent. We show that KneEZ Clear, which is based on the EZ Clear method, is highly flexible, demonstrating efficacy in a wide range of murine musculoskeletal tissues including the vertebral column, hindlimb, skull, and teeth. Critically, KneEZ Clear does not require specialized equipment and retains endogenous signal from fluorophores and fluorescent proteins. Additionally, following clearing and wholemount imaging, precious samples can still be processed for subsequent 2D histological analyses for validation or further study. Finally, we show that KneEZ Clear can be applied to samples of disease models to reveal alterations in tissue architecture and homeostasis. The simplicity, versatility, and efficiency of KneEZ Clear for optical clearing of musculoskeletal tissues will accelerate our understanding of cellular interactions and dynamics in homeostasis and disease.

## Introduction

Over the last 10-15 years, advances in three dimensional (3D)-imaging technologies have transformed biomedical research by enabling in-depth interrogation of the native, 3D architecture of tissue and entire organ systems with greater ease and accuracy than previously thought possible.^1^ Optical clearing methods, which minimize light-scatter or refraction within a sample, rendering it optically transparent to enable the visualization and analysis of entire organs and tissues in their native, 3D environment, have played an essential role in the evolution and revolution of 3D imaging.^1,2^

To understand how light interacts with a molecule or medium, researchers calculate its refractive index (RI), which reflects the ratio of the speed of light in a vacuum divided by the speed of light in the presence of the molecule or medium.^1^ To render a cell optically transparent, the RI of all light-scattering cellular components (nuclei, ribosomes mitochondria, lipid droplets, membranes, myelin, extracellular matrix components, and others) must match their surrounding environment, e.g., the imaging medium in which the sample is immersed. Beyond just cellular constituents, biological samples consist of connective tissue, cellular matrixes, and extracellular fluids, each with their own RI. For example, musculoskeletal tissues are comprised of both mineralized and non-mineralized constituents with varying amounts of lipids, collagen fibers, inorganic hydroxyapatite crystals, and hemoglobin that differentially affect light absorption and scattering, and produce background autofluorescence. These differences explain why bone has a greater RI (1.65-1.65) compared to cartilage (1.38-1.4), proteins and lipids (1.4-1.6), and water (1.33).^3,4^ Thus, to render tissues optically transparent, one must uniformly reduce these differences in light scatter throughout the sample, as this heterogeneity prevents the uniform passage of light through a sample.

Tissue clearing methods primarily fall into three categories: hydrophobic (solvent-based), hydrophilic (aqueous-based), and hydrogel-based methods.^3,5^ Generally, these methods rely on removing lipids (delipidation), pigment (bleaching), and can include calcium phosphate removal (demineralization) prior to RI matching of the specimen to the imaging media.^2^ Hydrophobic methods typically shrink tissues, while hydrogel- and hydrophilic-based methods (depending on the osmolarity of the reagents) can expand the specimen. While both hydrophobic- and hydrophilic-based methods are compatible with immunohistochemistry, hydrophilic-based clearing approaches show improved preservation of emission of endogenous fluorescent proteins. Additionally, in contrast to hydrogel-based methods, such as CLARITY, aqueous protocols do not require specialized equipment for sample processing^6,7^. Finally, aqueous-processed samples can typically be imaged at institutional imaging cores, unlike those cleared with organic-based clearing methods which often include corrosive or combustible reagents.^8^ Thus, aqueous clearing methods are highly preferred for their simplicity, flexibility, and ease-of-use.

Musculoskeletal tissues, such as bones, ligaments, muscles, tendons and teeth, are notoriously difficult to study using histological and imaging-based methods. These tissues typically have a complex extracellular matrix (ECM), contain high levels of endogenous fluorescence, and – in the case of bone – display age-dependent levels of calcification that complicate the sectioning and staining process. Imaging-based studies of physiological and pathological processes in these tissues have historically relied on 2-dimensional tissue processing workflows, which are labor- and time-intensive, and require serial sectioning to analyze the process-of-interest throughout the tissue.^9,10^ More recently, intravital multiphoton imaging emerged as a powerful technology to study cellular processes in skeletal tissues, but it has limited observation depth.^11,12^ Given the inherent three-dimensional structure of cells and organs, biologists are increasingly turning to volumetric imaging approaches, such as micro-CT and magnetic resonance imaging,^9,13^ though these two approaches have limited resolution compared to recently developed lightsheet fluorescent microscopes.^14–17^ Thus, there is an immense need for advanced tissue clearing methods that are adapted for musculoskeletal tissues and will enable 3D-imaging of cellular and tissue architecture and homeostasis across physiological and disease states.

Several effective clearing methods have been developed for musculoskeletal tissues, but all have critical limitations.^18–21^ Hydrogel-based methods (CLARITY, Bone CLARITY, PACT)^18,22,23^ are lengthy and typically require expensive, specialized equipment. Solvent-based methods (iDISCO, uDISCO, OsteoDISCO, WildDISCO, PEGASOS),^21,23–26^ are effective, but they may deform tissues, rapidly quench endogenous fluorescence, and require hazardous chemicals that are often not compatible with imaging platforms.^7^ A single aqueous-based method, CUBIC,^27^ has been developed for clearing intact mice, but it requires a lengthy, 2- to 3-week clearing process. Furthermore, most methods evaluate efficacy using isolated single tissues (e.g., bone or muscle)in order to reduce processing duration and increase clearing efficiency.^18–21^ Accordingly, the field would greatly benefit from a simple and fast, aqueous-based method for effective clearing of heterogeneous musculoskeletal tissues.

Here, we present KneEZ Clear, an simple, rapid, flexible, and reproducible technique for clearing murine skeletal tissues that is based on EZ Clear.^6,8^ By optimizing the 3 steps (fixation, delipidation, and RI-matching) of the original EZ Clear protocol, we show that KneEZ Clear generates optically-transparent adult mouse knee joints while retaining signal from endogenous fluorescent proteins. While initially developed for clearing mouse knee joints, KneEZ Clear efficiently renders other musculoskeletal tissues, such as the vertebral column, tooth, hindpaw, and skull optically transparent. Furthermore, we show that optically transparent wholemount adult murine knee joints that were processed with KneEZ Clear and imaged using lightsheet fluorescence microscopy (LSFM), can be further processed for cryosectioning, immunolabeling, and fluorescent imaging while retaining endogenous fluorescent signals. Finally, we applied KneEZ Clear on ankle joints of *FKBP10* mutant mice, a mouse model of osteogenesis imperfecta that develops a combined tendinopathy and osteoarthritis in multiple joints, and develops ankle joint contractures with age.^28^ Wholemount imaging of KneEZ Clear-processed *FKBP10* mutant ankle joints confirmed the presence of joint contractures. Importantly, it also revealed excessive vascularization at the site of contracture compared to control littermates. These results show that KneEZ Clear efficiently clears a variety of musculoskeletal tissues, and that it can be applied to study how biological tissues and processes change in the context of disease within intact joints.

## Material and Methods

### Mice

All mouse protocols were approved by the Institutional Animal Care and Use Committee (IACUC) at Baylor College of Medicine, the University of Virginia School of Medicine, and the University of Michigan School of Dentistry. For all experiments, the day of birth was considered P0, and all adult mice were at least 8 weeks of age. Mice were housed with access to food (normal chow diet) and water ad libitum on a 12-h light–12-h dark cycle at 21°C and 50–60% humidity.

*Prg4-BAC-CreERT2* mice (MGI ID: 7435587) were previously generated by Maurizio Pacifici,^29^ while Ai9 (*R26^CAG-lsl-TdTomato^*) mice (RRID: IMSR_JAX:017455) were purchased from Jax. Mice were genotyped by PCR using the following oligos: Fwd: 5’-TCC AAT TTA CTG ACC GTA CAC CAA-3’ Rev: 5’-CCT GAT CCT GGC AAT TTC GGC TA-3’, resulting in amplicons of ∼1,100 bp for the Prg4-Cre transgene, ∼480bp for *Rosa26* wildtype mice, and ∼240bp for the Ai9 allele. Tamoxifen (Sigma-Aldrich, #10540-29-1) solution was made at a concentration of 20 mg/mL by dissolving 200 mg of tamoxifen in 0.5 mL of 100% ethanol and subsequently adding sesame oil [Sigma-Aldrich, #8008-74-0] until a total volume of 10 mL. Adult *Prg4-BAC-CreER;R26^CAG-lsl-TdTom/CAG-lsl-TdTom^* mice were pulsed with 3 consecutive doses of tamoxifen (100 mg/kg body weight, injected intraperitoneally, once per day).

### Tissue collection and clearing

Perfusions and tissue collection were performed as described previously.^6^ Briefly, mice were deeply anesthetized by inhalation of 4-5% isoflurane and injected with 50 μL *Lycopersicon esculentum* (Tomato) Lectin DyLight 649 [Vector Laboratories, #DL-1178-1] via retroorbital perfusion, which was allowed to circulate for 15 minutes. After 15 minutes, animals re-anesthetized and opened to expose the thoracic cavity, then perfused with 100 μL lectin through the left ventricle. This was followed by perfusion with 5 mL warm (37°C) 1x PBS and then 5 mL room temperature (RT, 21°C) 1x PBS / 4% PFA. Finally, a post-fix pulse of 1 mL diluted lectin (1:10 in 1x PBS) was injected into the left ventricle. The tissue of interest (hind limb, ankle, knee joint, ankle, hind paw, spine) was dissected away, skin was removed manually, and muscles were optionally trimmed to isolate specific tissues of interest. Samples were then drop fixed in 1x PBS / 4% PFA overnight at a 1:20 volume:volume ratio to ensure adequate fixation and incubated at 4°C with gentle agitation, wrapped in foil to preserve fluorescent signal. The following day, samples were washed three times with 1x PBS at RT (21°C), on an orbital shaker with gentle agitation, one hour per wash. The samples were then immersed in 10% EDTA (1x PBS) for decalcification at RT for 2-5 days. X-ray imaging was used to confirm demineralization. For ankles, we found that extended demineralization (1 week) was required compared to knee joints. Samples were washed with 1x PBS (3X, 1hr/wash, RT) and then processed (dehydrated) through an increasing gradient of methanol in 1x PBS (50%, 80%, and 100%), 1 hour per wash, before incubation in a hydrogen peroxide solution (1 volume H_2_O_2_, 1 volume DMSO, 4 volumes 95% MeOH) overnight (16 hours) at 4°C to remove heme and decolorize the tissue. Samples were then washed in 100% methanol (2X, 1hr, RT), then taken through a reverse methanol:1x PBS gradient of 80% and 50% before going into 1x PBS. Following this, samples were taken through an increasing gradient of tetrahydrofuran (THF) [Sigma-Aldrich, #186562] with triethylamine (TEA) [Sigma-Aldrich, #471283] in ddH_2_O for 12 hours each: 50% THF (20μL TEA in 10ml THF + 9.98ml ddH_2_O), 70% THF (30μL TEA in 14ml THF + 5.97ml ddH_2_O), 80% THF (44.8μL TEA in 16ml THF + 3.95ml ddH_2_O), and 90% THF (64μL TEA in 18ml THF + 1.36 ml ddH_2_O) (x2). Samples were then washed in ddH_2_O (4X, 1hr/wash, RT) and then incubated in EZ View: 80% Nycodenz (Serumwerk Bernburg, #18003), 7 M urea, and 0.05% sodium azide. Samples were equilibrated in EZ View for 1-2 days until they were rendered optically transparent and then mounted in 1% filtered regular melt agarose (in ddH_2_O) with a sample holder for imaging. Once the agarose solidified, samples were re-equilibrated in EZ View for 1-2 days until transparent and EZ View media was changed prior to wholemount imaging on a Z.1 (Zeiss), custom-built Mesospim,^14,15^ or 3i CTLS (intelligent imaging) lightsheet microscope.

For clearing of murine molars: Animals were perfused with lectin and PFA as described above, prior to molar teeth extraction, with the removal of any excess bone tissue and were then drop fixed and washed as described for musculoskeletal tissues above. The samples were then immersed in 10% EDTA / 1x PBS for decalcification at RT for 6 days, refreshed every two days. Following decalcification, samples were washed with 1x PBS, one hour per wash, at RT and then taken through an increasing gradient of tetrahydrofuran (THF) [Sigma-Aldrich, #186562] ddH_2_O for 12 hours each: 50% THF, 70% THF, 80% THF, and 100% THF (x2). Samples were then washed four times, 1 hour per wash, in ddH_2_O and then incubated in EZ View, containing 80% Nycodenz (Serumwerk Bernburg, #18003), 7 M urea, and 0.05% sodium azide, for refractive index matching. Teeth were temporarily glued via a Bondic UV Glue kit to a bar to secure the sample, and equilibrated in a chamber filled with EZ view prior to imaging on the Zeiss Light Sheet 7. Images were taken at both 5X and 10X, at a refractive index value of 1.51.

A detailed version of the KneEZ Clear protocol is available on Protocols.io (dx.doi.org/10.17504/protocols.io.14egn3oxyl5d/v1).

### Immunofluorescence

KneEZ Clear-processed *Prg4^CreER^; R26^CAG-lsl-tdtomato^* knee joints were taken out of the EZ View RI-matching media and washed with 1x PBS (3X, 30min, RT) with gentle agitation. A final wash with 1x PBS was done for 1 hour. Following this, knees were further decalcified in 14% EDTA for three days with gentle agitation at 4°C. Samples were then washed with 1x PBS (3X, 30min, RT) with gentle agitation, and were cryopreserved in 30% sucrose in 1x PBS at 4°C for 24-48 hours, or until they sank to the bottom of the vial. Next, samples were embedded in Tissue-Tek^®^ O.C.T. (optimum cutting temperature; VWR; #25608-930) for coronal sections, using the dry ice block freezing method. 25 μm thick sections were stained with DAPI (FisherScientific, #NC0056218) solution for 30 minutes to label nuclei, then washed three times with 1X PBS, mounted with ProLong™ Glass Antifade Mountant (Thermo Fisher, #P36980), and cover-slipped with Precision Cover Glasses (FisherScientific, #NC1415511). For immunofluorescence staining, slides were stored in a dark slide chamber overnight at RT. Sections were fixed with 4% PFA for 30 minutes, followed by washes with 1x PBS. Samples were incubated with trypsin antigen retrieval solution (Abcam, Catalog #AB970) for 15 minutes. To reduce autofluorescence, sections were incubated for 1 hour in a working solution of prepared M.O.M. (Mouse on Mouse) Mouse IgG Blocking Reagent (Vector Laboratories, BMK-2202) in a dark slide chamber. Knee sections were incubated overnight in a humidified dark slide chamber with a primary antibody against Collagen II (mouse monoclonal antibody, ThermoFisher, Catalog #M2139) at a dilution of 1:50 in prepared M.O.M. Diluent (Vector Laboratories, BMK-2202), followed by secondary antibody incubation with anti-mouse Alexa Fluor 633 (Thermo Fisher, Catalog #A-21052) at a dilution of 1:200 for 1 hour. One drop of DAPI (Fisher Scientific, Catalog #NC0056218) was added for 30 minutes and then washed three times with 1× PBS. Samples were mounted with one drop of ProLong™ Glass Antifade Mountant (Thermo Fisher, Catalog #P36980) and cover-slipped using Precision Cover Glasses (Fisher Scientific, Catalog #NC1415511). Tile (4×4) images with a z-stack (3 μm, 6 slices) of knees were acquired using a Zeiss LSM 880 Airyscan FAST confocal microscope with a 10× objective

### Sample Mounting for Lightsheet Imaging

Cleared, RI-matched samples were mounted in filtered 1% agarose (in ddH_2_O) using a 3-5 mL syringe with the top cut off, positioned vertically over a test tube rack for stability. Prior to mounting, agarose was heated in a microwave and then vacuum filtered using a 0.45 μm bottle top filter. Filtered agarose was then cooled in a 37°C water bath. Each sample was individually placed in a syringe and warm (37°C) agarose was slowly added until the sample was covered with agarose by ∼1 cm. A sample holder screw was then embedded above the sample before the agarose solidified. Once the agarose solidified, samples were placed in EZ View RI matching media to re-equilibrate for 24-48h. On the day of imaging, a magnet is placed at the end of the embedded screw and used to attach the sample to the sample holder in the microscope.

### Lightsheet Image Acquisition and Processing

#### Zeiss Z1 Lightsheet Microscope

Mounted, RI-matched samples were imaged on a Zeiss Z1 Lightsheet Microscope. Samples were placed in the imaging chamber and submerged in EZ View refractive index matching media. The magnification was set to 5X. Image acquisition was done using a 640 nm laser set to 20% power, or a 561 nm laser set to 15% power, with an exposure time of 100 ms. Images were taken at an axial resolution of 1.8 μm and a Z-step of 3.6 μm in a tiled sequence with 20% overlap.

Acquired tiled datasets were stitched together using Stitchy software with the file output designated as an .ims file (Imaris compatible). Datasets were then visualized and manipulated in the Imaris software V10.2.0. Representative images of the reconstructed samples were captured using the Imaris snapshot function in the 3D view.

#### Cleared Tissue Lightsheet XL Microscope

Mounted, RI-matched samples were imaged on a 3i (Intelligent Imaging Innovations) Cleared Tissue Lightsheet XL Microscope (CTLS XL). Samples were placed in the imaging chamber and submerged in EZ View refractive index matching media. The magnification was set to 1X. Image acquisition was done using a 640 nm laser set to 200 mW, with an exposure time of 100 ms. Images were taken at an axial resolution of 0.8 μm and a Z-step of 11 μm in a tiled sequence with 15% overlap.

Acquired tiled datasets were stitched together using Slidebook software (3i). The datasets were converted to an .ims (Imaris compatible) file type using ImarisFileConverter V10.1.0. Datasets were then visualized and manipulated in the Imaris software V10.2.0. Representative images of the reconstructed samples were captured using the Imaris snapshot function in the 3D view. Digital sections were captured using the Imaris slice function at increasing imaging depth.

#### mesoSPIM Lightsheet Microscope

Mounted, RI-matched samples were imaged on a custom-built mesoSPIM Lightsheet Microscope. Samples were placed in the imaging chamber and submerged in EZ View refractive index matching media. The magnification was set to 2X. Image acquisition was done using a 561, 638, or 785 nm laser set to 15% (for 561 and 785 nm lasers) or 30% (for 638 nm laser). The exposure time was set to 20 ms. To capture reflected light images, the 785 nm laser was used without the emission filter. Images were taken at an axial resolution of 2.125 μm and a Z-step of 5 μm. Tiling was not done for these datasets.

Acquired datasets were converted to an .ims (Imaris compatible) file type using ImarisFileConverter V10.1.0. Datasets were then visualized and manipulated in the Imaris software V10.2.0. Representative images of the reconstructed samples were captured using the Imaris snapshot function in the 3D view. Digital sections were captured using the Imaris slice function at increasing imaging depth.

## Results

### Developing KneEZ Clear to efficiently clear musculoskeletal tissues

We recently developed EZ Clear as a simple, rapid, and robust method to clear whole murine organs.^8^ This aqueous-based clearing method can be applied in standard lab environments, but it was optimized for soft tissues. When we applied this method to the mouse hindlimb, the standard EZ Clear method displayed subpar efficiency at clearing musculoskeletal tissues (**Suppl Fig 1**). We therefore sought to optimize EZ Clear for clearing of murine mineralized tissues to accelerate studies of structural and cellular changes in musculoskeletal tissues in 3D.

We first focused our efforts on the murine knee joint, as it is composed of multiple tissues (bone, synovium, ligaments, fat pad, and others) that each have distinct mineral, lipid, and tissue stiffness profiles.^30^ To examine tissue clearing throughout the mouse hindlimb, we labeled the vasculature of 3-month-old wildtype (C57Bl/6J) mice by injecting them with a far-red fluorescently conjugated *Lycopersicon esculentum* (tomato) lectin (lectin-DyLight 649 nm) dye that binds endothelial cells.^31–33^ Following lectin-labeling and sample collection, we assessed perfusion and vascular labelling efficiency in the hindlimb and brain. To do so, samples were imaged using an epifluorescence microscope to visualize major vascular anatomical landmarks (**Fig 1A**). Importantly, we found that an additional transcardial pulse of lectin post-fixation (lectin chase) improved the perfusion and labeling efficiency of the vasculature in the hindlimb (**Suppl Fig 2**). Following confirmation of successful perfusion of large and small vessels, hindlimbs were subsequently processed with iterative modifications of the EZ Clear method to define optimal conditions for tissue fixation, decalcification, and delipidation of mineralized tissues **(Suppl Fig 3-A).**

**Figure 1:**
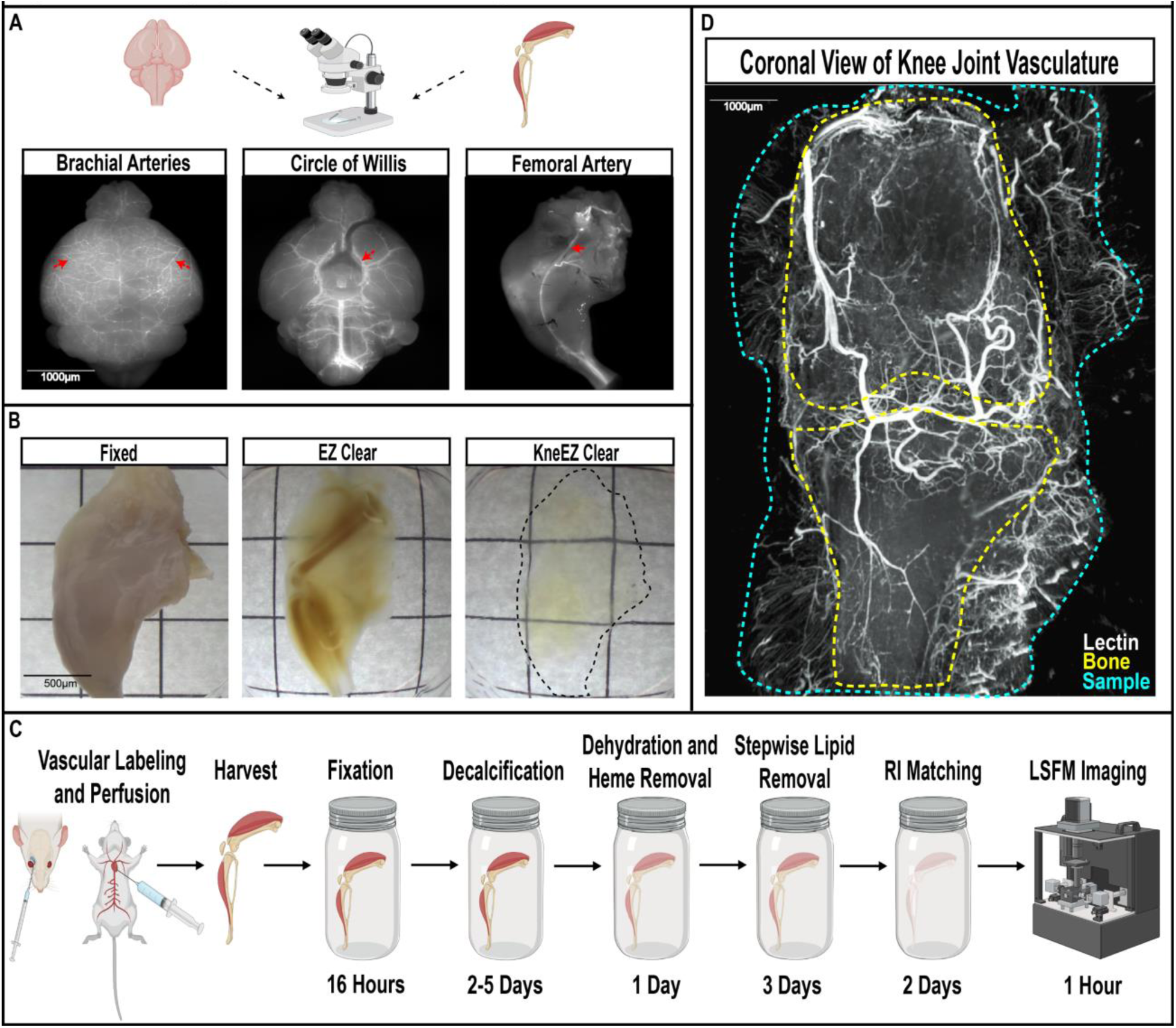
(A) Lectin-mediated fluorescence in fixed, uncleared brain and hindlimb samples, imaged with an epifluorescence microscope to verify successful lectin labeling. Vascular landmarks such as the brachial arteries (dorsal image of brain; left panel), circle of Willis (ventral image of brain; middle panel), or the femoral artery (medial view of hindlimb, right panel) marked with red arrows, should be visible. (B) Brightfield images of a fixed hindlimb (left), an EZ-Clear processed hindlimb (middle), and a KneEZ Clear-processed hindlimb (right) over grid paper reveals superior sample transparency of KneEZ Clear compared to EZ Clear. (C) Overview of the KneEZ Clear pipeline. A fluorophore-conjugated lectin is injected into the retro-orbital sinus and left ventricle to label the vasculature. This is followed by perfusion of PBS, 4%PFA, and a post-pulse of fluorophore-conjugated lectin prior to sample collection, which are then drop-fixed. Samples next undergo decalcification, heme removal, lipid removal, and refractive index matching to render samples transparent for imaging. (D) 3D reconstruction of a cleared knee joint (coronal view) imaged with a Zeiss Z1 Lightsheet microscope. Lectin-labeled vascular signal is shown in white, femur and tibia are outlined in yellow, and the sample is outlined in blue.

We focused on optimizing tissue clearing for mouse hindlimbs because they are composed of multiple joints, bones, muscles, ligaments, and tendons; reasoning that a method that efficiently clears entire hindlimbs would work on other musculoskeletal tissues. First, we altered the fixation and decalcification parameters, varying the duration (16, 24, or 48 hours for fixation; 2 days or 2 weeks for decalcification), temperature (4°C or 21°C), and concentration (10 or 14% EDTA) of incubation steps. We chose to focus on using EDTA-based demineralization because it preserves tissue organization, antigen activity, and fluorescent reporter signal.^34^ We found that a 16-hour fixation at 4°C and a 2-day decalcification at 21°C produced minimal autofluorescence (**Suppl Fig 3-B**).

Next, we varied decolorization parameters to remove heme-based pigments and further reduce tissue autofluorescence. We compared hydrogen peroxide (H_2_O_2_)- and amino-alcohol-mediated bleaching methods as they were previously shown to be effective for tissue clearing.^18,24^ Stepwise dehydration and bleaching with H_2_O_2_ and methanol, as described by Renier et al.,^24^ yielded the best results for murine hindlimbs (**Suppl Fig 3-B**).

Finally, we modified EZ Clear’s delipidation step, which consists of an overnight incubation in 50% tetrahydrofuran (THF).^8^ We found that stepwise delipidation (50%-70%-80%-90% in ddH_2_O, 12h each at 21°C) with the water-miscible tetrahydrofuran (THF), as described by Erturk et al.,^35^ improved lectin’s signal to noise ratio in mouse hindlimbs compared to a single 16-hour incubation step in 50% THF (**Suppl Fig 3-B**).^36^ We then tested the effect of adding triethylamine (TEA) to each THF-incubation step, increasing the solution’s pH which should preserve endogenous fluorophores.^35^ We included this step to preserve native, endogenous reporters and saw no negative impact on lectin dye labeling in the mouse hindlimb.

In sum, we found that overnight fixation at 4C and 2-day decalcification, followed by a stepwise delipidation through THF in the presence of TEA improved EZ Clear-based optical clearing of the entire knee joint (**Suppl Fig 4**). We named this mineralized tissue version of EZ Clear, KneEZ Clear.

We next assessed and compared the original EZ Clear and the optimized KneEZ Clear methods’ ability to render murine hindlimbs optically transparent. Wildtype murine hindlimb samples processed using the KneEZ Clear protocol achieved optical transparency throughout the entire joint, as light transmission through the sample was observed using both brightfield (**Fig 1B**) or LSFM imaging (**Fig 1D**). In contrast, brightfield and LSFM imaging of EZ Clear-processed revealed incomplete light transmission and elevated autofluorescence levels throughout the sample (**Fig 1B; Suppl Fig 1**). In addition, optical sections at multiple positions along the sample’s anteroposterior axis revealed higher signal-to-noise ratios for lectin-labeled vasculature at all imaging depths for KneEZ Clear’ed samples compared to EZ Clear’ed samples (**Suppl Fig 1**). Together, these data confirm KneEZ Clear has superior performance for rendering mouse hindlimbs optically transparent.

KneEZ Clear effectively renders the adult mouse hindlimb optically transparent in 10 days using 5 simple steps (**Fig 1C**). Following perfusion and dissection, hindlimbs are immersion-fixed overnight in 4% PFA (<u>Step 1</u>), then decalcified for 2 days using 10% EDTA (<u>Step 2</u>). Samples are then dehydrated using a methanol gradient and bleached in hydrogen peroxide to remove the remaining heme present in the muscle and bone marrow (<u>Step 3</u>). Next, hindlimb samples undergo a series of lipid removal steps using increasing concentrations of tetrahydrofuran (<u>Step 4</u>) in TEA. To achieve optical transparency, the hindlimb samples are then put in EZ View,^8^ an aqueous RI-matching solution that allows greater depth of imaging than other aqueous solutions such as RIMS (<u>Step 5</u>).^8^ Once samples are fully equilibrated in this RI matching solution, they are ready for imaging.

### KneEZ Clear efficiently clears the murine knee joint and preserves endogenous fluorescence

LSFM imaging of lectin-perfused, KneEZ Clear-processed wildtype mouse hindlimbs to visualize lectin-labeled (656-720 nm) vasculature structures revealed a dense network of blood vessels that permeates the sample, including the popliteal artery that branches into the superior and inferior genicular arteries around the knee joint, and the saphenous artery that descends down the medial side of the hindlimb towards the ankle (**Fig 2-A, Suppl Video 1**). Virtual sections from the 3D reconstruction showed well-defined vascular signal throughout the mediolateral axis when imaged in the coronal plane, regardless of imaging depth (**Fig 2-B**), showing effective clearing and imaging throughout the knee joint.

**Figure 2:**
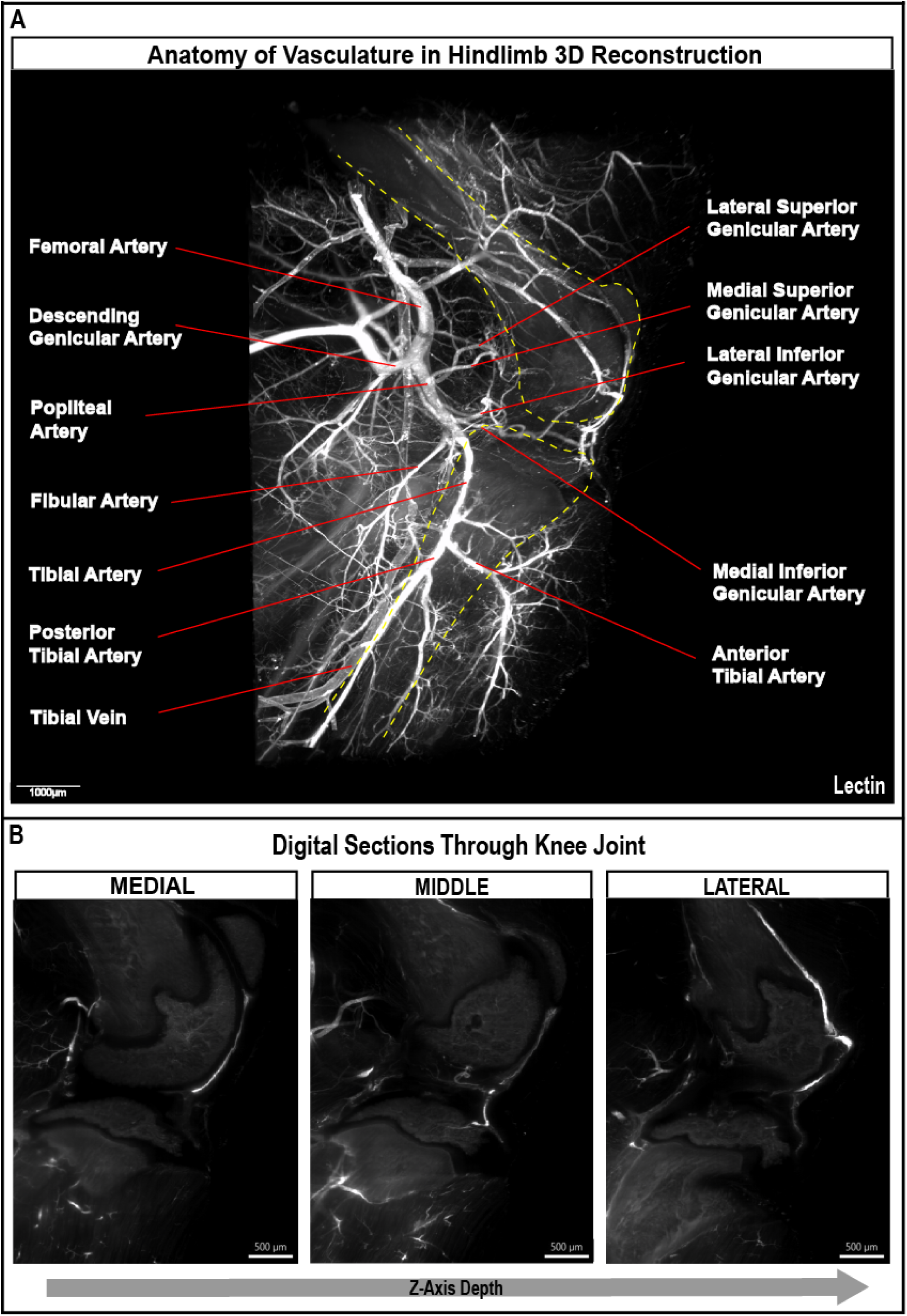
(A) 3D reconstruction of a KneEZ Clear’ed, lectin-perfused hindlimb (sagittal view, medial side), imaged using 3i’s AxL Cleared Tissue Lightsheet. The vascular network (white; lectin) throughout the knee joint and associated tissues is visible, including major anatomical vessel landmarks such as the femoral and genicular arteries. Femur and tibia are outlined in yellow. (B) Digital sections throughout the knee joint reveal sharply outlined lectin signal throughout the knee joint, from the medial side (closest to the camera) to the lateral side (furthest away from the camera).

A major advantage of aqueous tissue clearing methods in general, and EZ Clear specifically, is the preservation of endogenous protein fluorescence.^8^ To validate that our optimized KneEZ Clear processing pipeline retained emission of endogenous fluorescent proteins, we cleared hindlimbs from *Tg(Prg4-BAC-CreER): R26^TdTomato/TdTomato^* mice. In these animals, expression of the Cre-dependent red fluorescent protein reporter, tdTomato, requires the activity of a tamoxifen-inducible Cre recombinase fused to a mutant estrogen receptor (CreERT2) inserted at the start codon of the *Proteoglycan 4* locus in a randomly integrated bacterial artificial chromosome (BAC) transgenic line.^29,37^ Prg4 (Proteoglycan 4, also known as lubricin) encodes a chondroprotective glycoprotein that is expressed in superficial zone chondrocytes and synoviocytes and plays a role as a boundary lubricant in articulating joints.^38,39^ Importantly, the *Tg(Prg4-BAC-CreER)* transgenic line recapitulates endogenous *Prg4* expression, as tamoxifen injection leads to CreERT2 recombination in articular cartilage, cruciate ligaments, and synovium.^29^ Mesospim-based imaging of KneEZ Cleared knee joints of tamoxifen-induced *Tg(Prg4-BAC-CreER): R26^TdTomato/TdTomato^* mice revealed robust tdTomato signal in the articular cartilage, synovium, and cruciate ligaments throughout the sample, in agreement with previous reports (**Fig 3A-C, Suppl Video 2**).^29^ While *Tg(Prg4-BAC-CreER): R26^TdTomato/TdTomato^* animals that did not receive tamoxifen displayed fluorescence in the same tissues (**Suppl Fig 5-B**), it was at a substantially lower intensity compared to induced samples (**Suppl Fig 5-C**). In addition, this fluorescent signal was absent in samples from tamoxifen-induced *Tg(Prg4-BAC-CreER)* animals that did not contain the *R26^TdTomato/TdTomato^* allele (**Suppl Fig 5-A**), suggesting low-level Cre recombination from the *Prg4^CreER^* allele under standard conditions. Importantly, sample excitation at 875 nm to collect reflective light from these samples enabled us to visualize gross anatomical features of knee-joint associated tissues, including tendons, ligaments, and even the tibial and femoral growth plates (**Fig 3C**).

**Figure 3:**
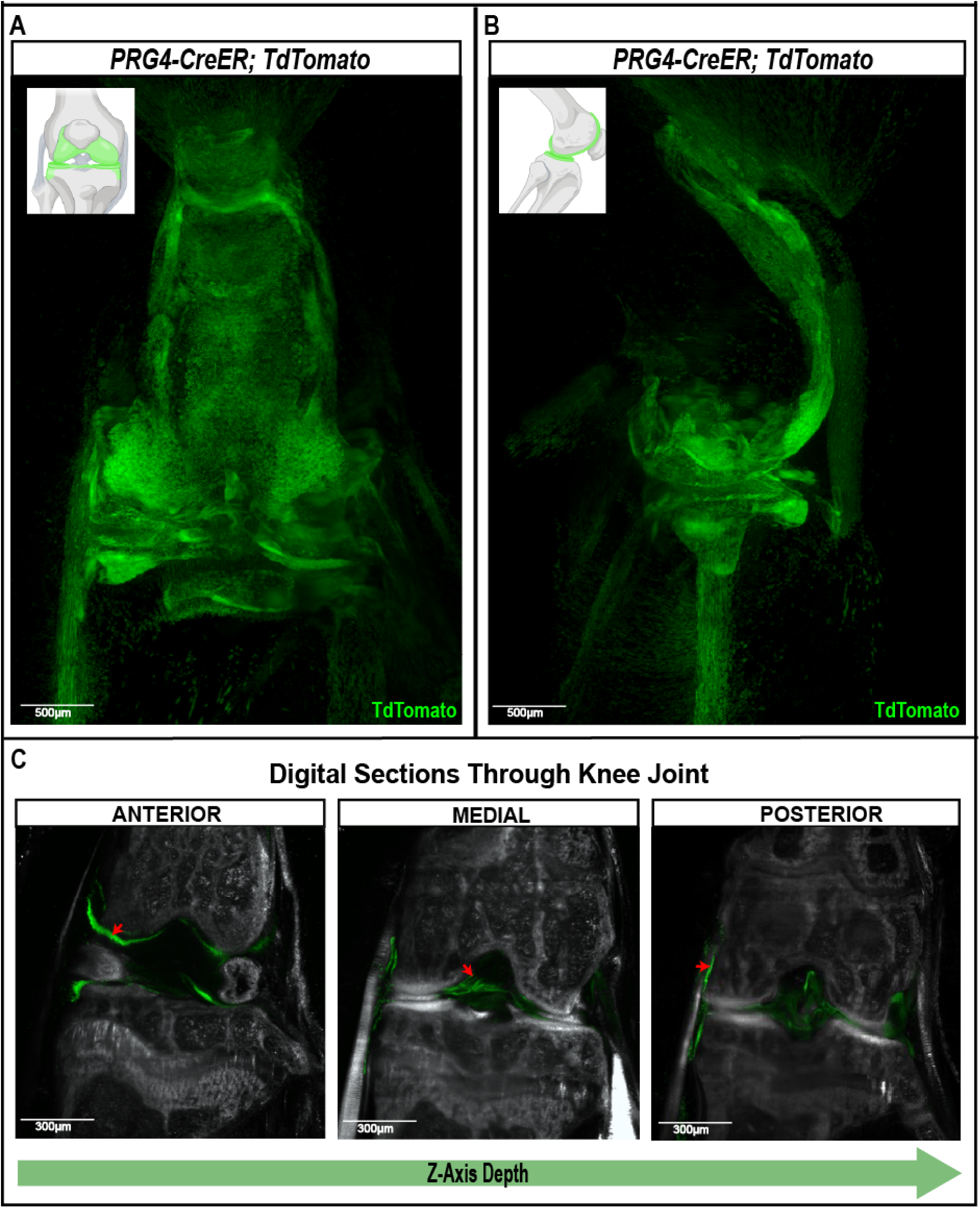
(A-B) Knee joints from *Tg(Prg4-BAC-CreER): R26^TdTomato/TdTomato^* mice in the coronal and sagittal view, showing preservation of Cre-induced fluorescent signal (green) in KneEZ Clear’ed samples. The expected expression pattern is displayed in the models in the top left corner of the images. (C) Digital sections throughout the knee joint (coronal view), with z-depth increasing from anterior to posterior revealing Cre-induced tdtomato expression in articular cartilage, cruciate ligaments, and synovium (red arrows in left, middle, and right panels respectively) as identified by reflective light (grayscale) collected from the sample, imaged with a MesoSPIM.

The exposure time was set to 20 ms. To capture reflected light images, the 785 nm laser was used without the emission filter

Previous reports showed that EZ Clear is compatible with cryosection-based histology and immunofluorescence staining following tissue clearing and wholemount imaging.^8^ To verify the fidelity of the observed *PRG4* lineage contribution revealed by our 3D imaging analysis patterns, we processed the KneEZ Clear-processed and 3D-imaged knee joints from tamoxifen-induced *Tg(Prg4-BAC-CreER): R26^TdTomato/TdTomato^* and *Tg(Prg4-BAC-CreER)* animals, as well as and those from non-induced *Tg(Prg4-BAC-CreER): R26^TdTomato/TdTomato^* animals using 2D histology and immunofluorescence. Confocal imaging of thick (25 µm) cryosections, stained with DAPI to visualize nuclei, confirmed robust tdTomato-expression in the articular cartilage, cruciate ligaments, and synovium of tamoxifen-induced samples (**Suppl Fig 5**). We next tested if KneEZ Cleared samples could be used for immunohistochemistry. Given the expression pattern we observed in *Tg(Prg4-BAC-CreER): R26^TdTomato/TdTomato^* knees, we stained cryosections from cleared knees for collagen II, which is expressed in articular and growth plate chondrocytes, and has also been detected in meniscal fibrochondrocytes.^40,41^ Confocal imaging of stained slides revealed significant overlap between tdTomato- and collagen 2-expressing cells in the articular cartilage and menisci of tamoxifen-induced *g(Prg4-BAC-CreER): R26^TdTomato/TdTomato^* animals (**Suppl Fig 6-C**) that was absent from uninduced animals (**Suppl Fig 6-B**), or animals that lacked the tdTomato allele (**Suppl Fig 6-A**). In sum, KneEZ Cleared samples that have undergone wholemount imaging can be processed for downstream cryosection-based histology and immunolabeling to confirm or expand upon findings.

### Reproducible, efficient clearing of other musculoskeletal tissues with KneEZ Clear

As noted, mouse hindlimbs are composed of multiple joints, bones, muscles, ligaments, and tendons. Thus, we reasoned that KneEZ Clear should be effective for whole mount clearing of other musculoskeletal tissues. To this end, we collected ankles, spines, and skulls from *Prg4^CreER^; ai9* animals. Similar to the murine knee joint, KneEZ Clear rendered all other musculoskeletal tissues optically transparent and permitted light transmission throughout the sample (Fig 4).^14,15^ In addition, mesospim-based imaging revealed intricate *Prg4-BAC-CreER* activity in facet joints of the adult murine vertebral column (**Fig 4-A**), metatarsophalangeal and interphalangeal joints in the hind paw (**Fig 4-B**), the skull, including the temporomandibular joint (**Fig 4-C**), and all joints that make up the ankle (**Fig 4-D**), confirming the application of this method for imaging other musculoskeletal tissues.

**Figure 4:**
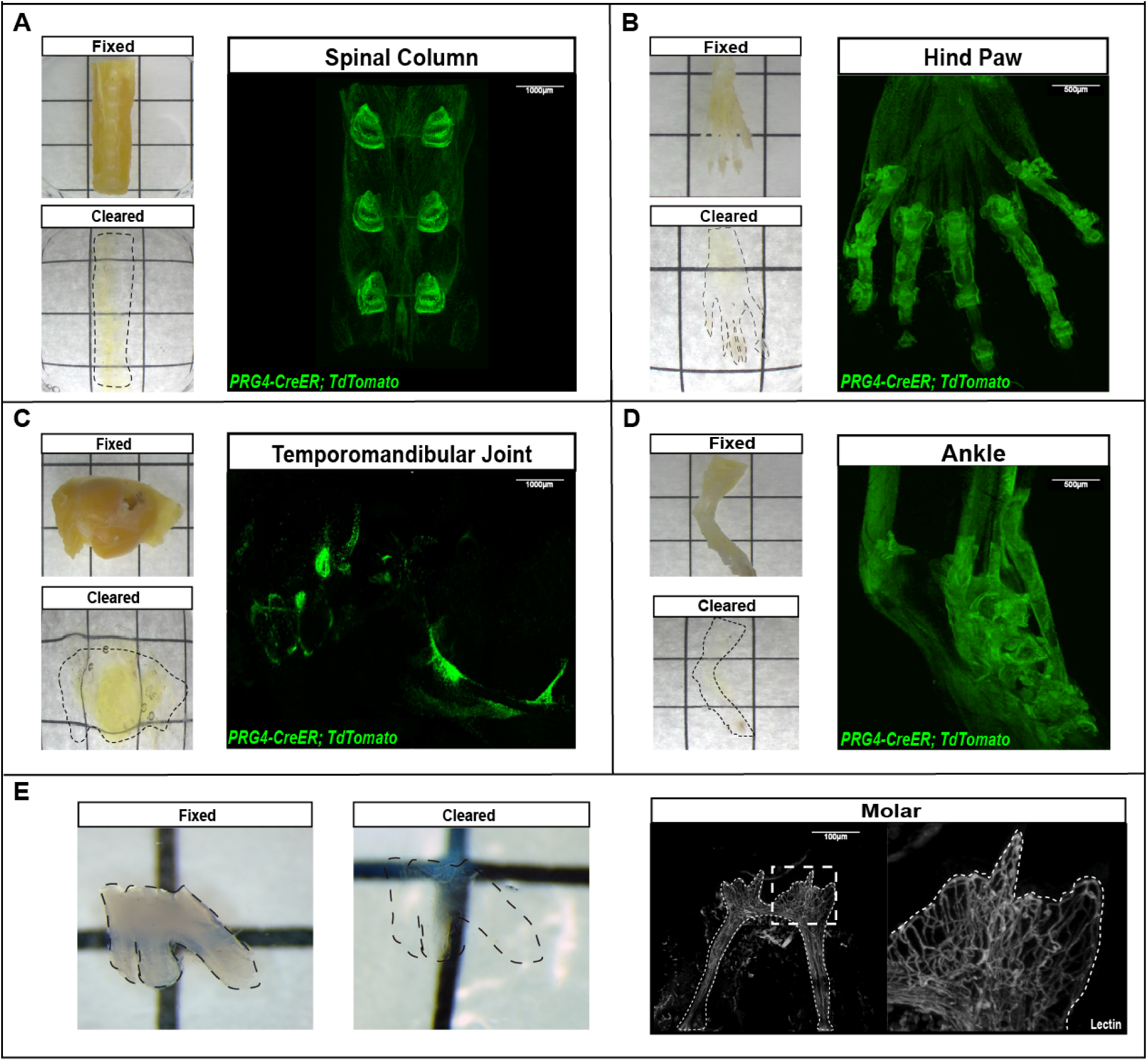
(A-D) Brightfield images of uncleared (fixed) and cleared musculoskeletal tissues obtained from *Tg(Prg4-BAC-CreER): R26^TdTomato/TdTomato^* mice, as well as 3D-reconstructed images of PRG-4-induced tdtomato-expression (green) in a KneEZ-Clear’ed spinal column (A), hind paw (B), temporomandibular joint (C), and ankle (D), imaged with a MesoSPIM. (E) Brightfield images of an uncleared (fixed) and cleared M1 molar, extracted from a wildtype mouse (left). 3D reconstructed images of KneEZ-Clear’ed molar, imaged with a Zeiss Light Sheet Z7 shows lectin-labeled vascular signal (white) within the inner dental pulp of the tooth.

We further tested KneEZ Clear’s ability to clear other complex hard tissues, such as mouse teeth, which consist of dense layers of dentin and enamel encapsulating and protecting the highly vascularized and innervated dental pulp.^42^ To this end, we perfused wildtype mice with lectin-DyLight 649 as described above, extracted M1 molars and submitted them to the KneEZ Clear protocol. Importantly, we found that an extended decalcification step (6 compared to 2 days) was needed to render this tissue optically transparent, as evidenced by brightfield and LSFM imaging (**Fig 4-E**). LSFM imaging showed strong fluorescent signal throughout the tooth, revealing the vasculature of the entire tissue (**Fig 4-E**). In sum, KneEZ Clear efficiently and reproducibly clears various types of heterogenous musculoskeletal tissues, including knee joints, hindpaws, vertebral columns, skulls, and teeth.

### KneEZ Clear permits the study of disease-induced vascularization changes in musculoskeletal tissues

The ultimate goal of tissue clearing and 3D imaging approaches is to reveal changes in tissue homeostasis and biological processes that are induced by age, disease, treatment, or other biologically relevant factors, and to study the mechanisms that drive them. We hence set out to determine if wholemount imaging of KneEZ Cleared samples yields the optical transparency and resolution needed to reveal such biologically relevant changes in musculoskeletal tissues. To do so, we cleared ankle joints of *Scx-Cre; FKBP10^fl/fl^* animals, a genetic mouse model for studying the pathogenic mechanisms underlying joint contractures in patients with osteogenesis imperfecta and Bruck syndrome.^28^ FKBP65, encoded by the FK506 Binding Protein 10 *(FKBP10) gene,* functions as a chaperone for type I procollagen and elastin, and its conditional loss in tendons and ligaments, and chondrogenic precursors (driven by *Scx-Cre*-mediated recombination) causes tendinopathy and osteoarthritis resulting in heterotopic ossification with aging.^28^ Wholemount imaging of KneEZ Cleared ankle joints of 8- to 12-week-old *Scx-Cre; FKBP10^fl/fl^* with our custom-built Mesospim instrument confirmed previously reported ankle joint contractures compared to controls (**Suppl Fig 7**). Importantly, assessing vascular networks in an additional set of 8- to 12-week-old *Scx-Cre; FKBP10^fl/fl^* ankle joints that were lectin-labeled revealed major vascular remodeling, with a strong increase in blood vessels at the site of contracture in mutants compared to *FKBP10^fl/fl^* controls (**Fig 5**). These data show that KneEZ Clear can be applied to intact joints to study how biological tissues and processes change in the context of complex joint disease.

**Figure 5:**
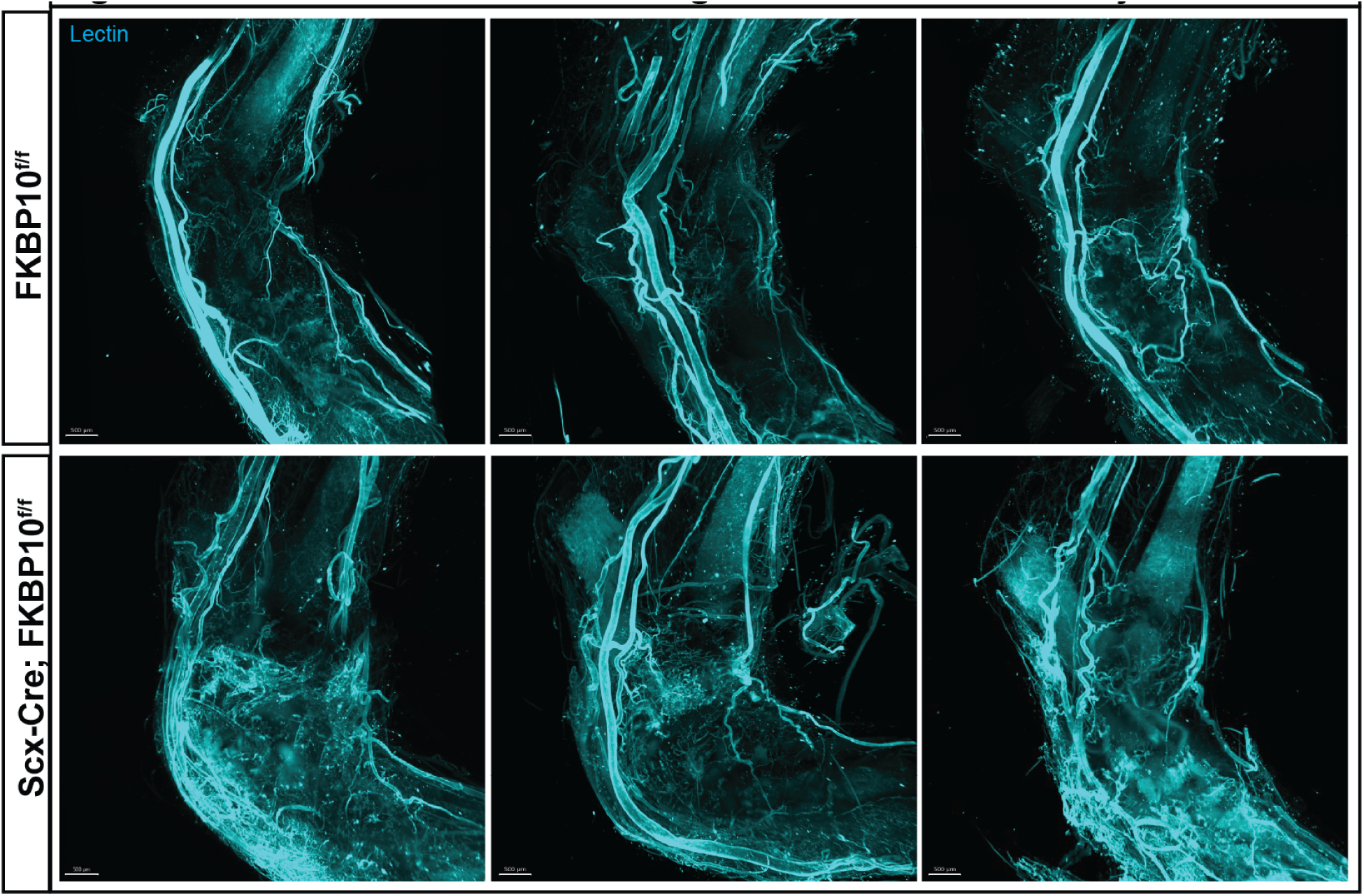
3D-reconstructed images of KneEZ Clear’ed, intact ankle joints from *FKBP10 ^f/f^* control (top) and *Scx-Cre; FKBP10 ^f/f^* (bottom) mutant animals, imaged with a Zeiss Z1 Lightsheet microscope. Lectin-labeled blood vessels (cyan) reveal increased vascularization at the site of joint contracture in *FKBP10* mutant animals compared to controls.

## Discussion

We have developed a simple, robust tissue clearing method to efficiently clear complex, heterogeneous musculoskeletal tissues. KneEZ Clear is based on EZ Clear, an aqueous-based tissue clearing method that does not use any special equipment or toxic organic solvents, making it easy to adopt and adapt to a research project’s imaging needs (**Fig 1**).^8^

KneEZ Clear can be used to visualize microanatomical structures within mineralized and non-mineralized musculoskeletal tissues. Here, we demonstrated this method’s ability to preserve fluorescent signals from fluorophores and fluorescent proteins in two ways. First, we visualized hindlimb vascularization by injecting animals with fluorescently conjugated lectin. Wholemount LSFM imaging of KneEZ Clear’ed hindlimbs revealed the extensive vascular network that permeates the muscles and joints of the mouse (**Fig 2**). Second, we imaged joints from *Tg(Prg4-BAC-CreER): R26^TdTomato/TdTomato^* mice, to visualize the previously Cre-activity of this line in articular cartilage, synovium, and cruciate ligaments in the knee joint (**Fig 3,4**). ^38,39^ In addition, we showed that cleared knee joints that had undergone wholemount imaging can be processed for downstream cryosectioning and immunohistochemistry to follow up on 3D based findings in 2D. Therefore, the aqueous-based solutions used for KneEZ Clear-based clearing of musculoskeletal tissues preserves fluorescence, unlike solvent-based solutions, such as BABB and DBE, which modify emission spectra of fluorophores and quench it over time.^43^ In addition, the imaging solution used for mounting and imaging KneEZ Clear samples, EZ View, has a high RI (1.518) and is not corrosive to microscopes, broadening its applicability and ease-of-use.^8^ Furthermore, because KneEZ Clear uses aqueous-based solutions at all steps of the protocol, processed samples should not expand or shrink. In contrast, hydrophobic (iDISCO, PEGASOS, BoneClear)^19,24,44^ or hydrogel-based (CLARITY, Bone CLARITY)^18,23^ methods developed for clearing musculoskeletal tissues can deform tissues.^7,8^ Therefore, aqueous-based tissue clearing methods such as KneEZ Clear or EZ Clear may enable more reliable downstream cross-comparisons or registration to a common sample due to the absence of tissue formation. Finally, we show that KneEZ Clear effectively renders other mineralized tissues optically transparent, including mouse spines, skulls, ankles, teeth, and hind paws (**Fig 4**).

Importantly, we show that KneEZ Clear-processed and samples that have undergone wholemount imaging can be processed for downstream cryosection-based histology and immunolabeling (**Suppl Fig 5, 6**). This is an important benefit of EZ Clear-based clearing methods that, to our knowledge, does not apply to other clearing methods. The ability to process samples after imaging for immunohistochemistry-based analysis allows researchers to assess additional proteins-of-interest and confirm or expand upon findings. We observed substantial consistency of fluorescent signal in similar tissue types within the knee using 2D and 3D imaging methods. However, 3D imaging revealed *PRG4-BAC-CreER* activity in anterior and posterior areas of the joint, areas that are typically bypassed for 2D histology in favor of sections in the middle area of the joint where both cruciate ligaments are visible (**Fig 3A, Suppl Video 2, Suppl Fig 5**). Hence, volumetric and complete imaging of an entire tissue by 3D wholemount imaging captures all information and provides deeper, structural insights compared to 2D imaging. However, 2D and 3D imaging methods can be used in a complementary fashion for KneEZ Cleared samples to confirm observed findings, e.g. cellular identities, or explore novel directions.

As shown here, optimization will be required for clearing each novel mineralized tissue in terms of either fixation (duration, temperature) decalcification (duration, temperature), or delipidation parameters in terms of defining their consequences on autofluorescence, tissue deformation, and sample transparency. For example, we found that extended fixation of murine joints increased autofluorescence in the mouse femur and tibia (**Supp Fig 3-B**). Of note, we found that the highly mineralized mouse tooth necessitated additional decalcification. Thus, we predict that decalcification parameters will also need to be modified based on the animal’s age and mineral content. For instance, minor adjustments to fixation and decalcification steps are needed for other skeletal tissues or other timepoints

Lastly, we showed, for the first time, that *FKBP10* mutant ankle joints display excessive vascularization that is associated with tendinopathy, osteoarthritis, and accelerated chondro-ossification, causing joint contractures (**Suppl Fig 6, Fig 5**). Both joint contractures, which were previously reported by Lim et al.,^28^ as well as previously unreported vascular alterations could be observed in KneEZ Cleared samples, showcasing KneEZ Clear’s ability to reveal biologically relevant tissue microarchitecture changes in whole joints. It remains unknown whether the excessive vascularization observed in *FKBP10* mutants precedes chondro-ossification of the tendons and ligaments in the ankle joint, and whether it plays a causative role in this process. Interestingly, our previous study identified upregulation of several molecules involved in vascular remodeling in *FKBP10* mutant ankle joints, including *Snai1*, *Tnn*, and *Serpine1*.^28^ Future studies will be needed to explore the cellular and molecular mechanisms that underlie the vascular phenotype in these mutants, and to reveal their connection to the documented changes in joint and motor function.^28^

In sum, KneEZ Clear is an easily adoptable, reproducible tissue clearing method that renders a wide variety of musculoskeletal tissues optically transparent while retaining fluorescent signals from labeled cell types of interest. Cleared samples can be further processed for downstream analyses, including standard histological analysis or immunohistochemistry and confocal imaging, accelerating the study of physiological and pathological processes in tissues that are notoriously difficult to image and study.

## Supporting information

Supplemental Figures_Combined

## Author Contributions

NAH, JDW, and BHL conceptualized the study. JY and NAH wrote the original draft. JY, LT, JDW, and NAH were involved in the design of experiments. JY, TA, LT, AIO, CL, SKP, and AG executed, imaged, and analyzed experiments. JMK performed mesospim image acquisition. All authors revised the paper and consented to its contents.

## Acknowledgements

We thank Dr. Maurizio Pacifici for sharing the genetic mouse line *Tg(Prg4-BAC-CreER)* with us. We are grateful to members of the RE-JOIN consortium for their suggestions on this manuscript. This project was supported by the Optical Imaging and Vital Microscopy (OiVM) Core at Baylor College of Medicine, by the University of Virginia Comprehensive Cancer Center and Intelligent Imaging Innovations, Inc (3i, Denver, CO, USA), and by the University of Michigan. The authors are also grateful to members of the RE-JOIN Consortium for their insights and suggestions on clearing knee joints and other musculoskeletal tissues. This work was supported by grants from the National Institute of Arthritis and Musculoskeletal and Skin Diseases of the National Institutes of Health through the NIH HEAL Initiative (https://heal.nih.gov/) under award number UC2AR082200 to B.L. and J.D.W. and UC2AR082197 to J.E. The content is solely the responsibility of the authors and does not represent official views of the National Institutes of Health.

The RE-JOIN consortium consists of The RE-JOIN consortium consists of Taeyoung (Ted) Ahn, Armen Akopian, Kyle Allen, Alejandro Almarza, Benjamin Arenkiel, Maryam Aslam, Basak Ayaz, Yangjin Bae, Bruna Balbino de Paula, Anita Bandrowski, Mario Danilo Boada, Jacqueline Boccanfuso, Jyl Boline, Dawen Cai, Dellina Lane Carpio, Robert Caudle, Racel Cela, Yong Chen, Rui Chen, Brian Constantinescu, Ibdanelo Cortez, Yenisel Cruz-Almeida, M. Franklin Dolwick, Chris Donnelly, Zelong Dou, Joshua J. Emrick, Malin Ernberg, Danielle Freburg-Hoffmeister, Jeremy Friedman, Spencer Fullam, Janak Gaire, Akash Gandhi, Terese Geraghty, Benjamin Goolsby, Stacey Greene, Nele Haelterman, Zhiguang Huo, Michael Iadarola, Shingo Ishihara, Azeez Ishola, Sudhish Jayachandran, Zixue Jin, Alisa Johnson, Frank Ko, Priya Kulkarni, Zhao Lai, Brendan Lee, Yona Levites, Carolina Leynes, Jun Li, Martin Lotz, Lindsey Macpherson, Tristan Maerz, Camilla Majano, Anne-Marie Malfait, Maryann Martone, Simon Mears, Bella Mehta, Emilie Miley, Rachel Miller, Richard Miller, Michael Newton, Alia Obeidat, Soo Oh, Merissa Olmer, Dana Orange, Miguel Otero, Kevin Otto, Folly Patterson, Marlena Pela, Daniel Perez, Sienna Perry, Theodore Price, Hernan Prieto, Russell Ray, Dongjun Ren, Margarete Ribeiro Dasilva, Alexus Roberts, Elizabeth Ronan, Oscar Ruiz, Shad Smith, Mairobys Soccorro Gonzalez, Kaitlin Southern, Joshua Stover, Michael Strinden, Hannah Swahn, Evelyne Tantry, Sue Tappan, Luis Tovias Sanchez, Cristal Villalba Silva, Airam Vivanco-Estella, Robin Vroman, Joost Wagenaar, Lai Wang, Kim Worley, Joshua Wythe, Jiansen Yan, and Julia Younis.

---

<u>Supplemental Figure 1:</u>

(A) 3D reconstruction of lectin-labeled (white) knee joints displaying the coronal (left) and sagittal view (right) of EZ Clear’ed (Top) or KnEZ Clear’ed (bottom) adult mouse hindlimbs, imaged with Zeiss Z7 Lightsheet microscope. (B) Digital sections throughout the knee joints of samples shown in A (coronal view), with z-depth increasing from anterior to posterior.

<u>Supplemental Figure 2</u>

(Left) An overview of the process of vascular labeling and perfusion comparing the original vascular labeling protocol, in which samples are immediately dissected after PFA perfusion, to an updated vascular labeling protocol that includes a post-fixation pulse with fluorophore-conjugated lectin prior to dissection (left). (Right) Brightfield images of brains and hindlimbs following the original vascular labeling protocol (top), and the updated protocol (bottom), showing improved vascular labelling efficiency by including an additional injection with labelled lectin.

<u>Supplemental Figure 3:</u>

(A) Table displaying the various parameters that were altered and tested for their impact on clearing efficiency. (B) 3D reconstruction of cleared knee joints (coronal view) displaying parameters that resulted in decreased clearing efficiency compared to other conditions. Cleared knee joints were imaged with a Zeiss Z1 Lightsheet microscope

<u>Supplemental Figure 4:</u>

(A) Comparison of EZ Clear and KneEZ Clear protocols, displaying differences during lipid removal (single step vs gradient THF-mediated delipidation), and the addition of triethylamine to the THF solution and an additional heme removal step for the KneEZ Clear method compared to the original EZ Clear protocol.

<u>Supplemental Figure 5:</u>

(A-C) Confocal images of Dapi-stained, coronal cryosections of knees from tamoxifen-induced, KneEZ Clear’ed, and 3D imaged *PRG4-CreER* (A) *or PRG4-CreER; TdTomato* (C) mice, and of uninduced *PRG4-CreER; TdTomato* mice (B). Top: Merged image showing dapi-stained nuclei (blue), and TdTomato (red), reflecting cells with current or previous PRG4-activity. Bottom: single channel (TdTomato) in greyscale.

<u>Supplemental Figure 6:</u>

(A-C) Confocal images of coronal cryosections of knee joints from tamoxifen-induced, KneEZ-Cleared and 3D-imaged *PRG4-CreER* (A) *or PRG4-CreER; TdTomato* (C) mice, and of uninduced *PRG4-CreER; TdTomato* mice (B) that were immunolabeled for collagen 2 (COLII). Top: Merged image showing COLII staining (green), dapi-stained nuclei (blue), and TdTomato (red), reflecting cells with current or previous PRG4-activity. Middle: single channel (TdTomato) in greyscale. Bottom: single channel COLII antibody staining in greyscale.

<u>Supplemental Figure 7:</u>

KneEZ-Clear’ed ankle joints from *Scx-Cre; FKBP10 ^f/f^* mice (8 weeks old), imaged on a MesoSPIM to collect light reflected off the sample. The reflected light captures signal from both bone and soft tissue, revealing joint contractures in *Scx-Cre; FKBP10 ^f/f^* ankles (B) that are not present in control littermates (A). A magnified view of the calcaneous is displayed in both panels, where the contracted tendon and reduced angle of the calcaneous relative to the tibia are visible.

