## Supplemental Figures_Combined for "KneEZ Clear, an Effective Tissue Clearing Protocol to Study Musculoskeletal Tissues in the Mouse"

Supplemental Figure 1. Comparing KneEZ Clear and EZClear's Efficiency at Clearing Mouse Knees

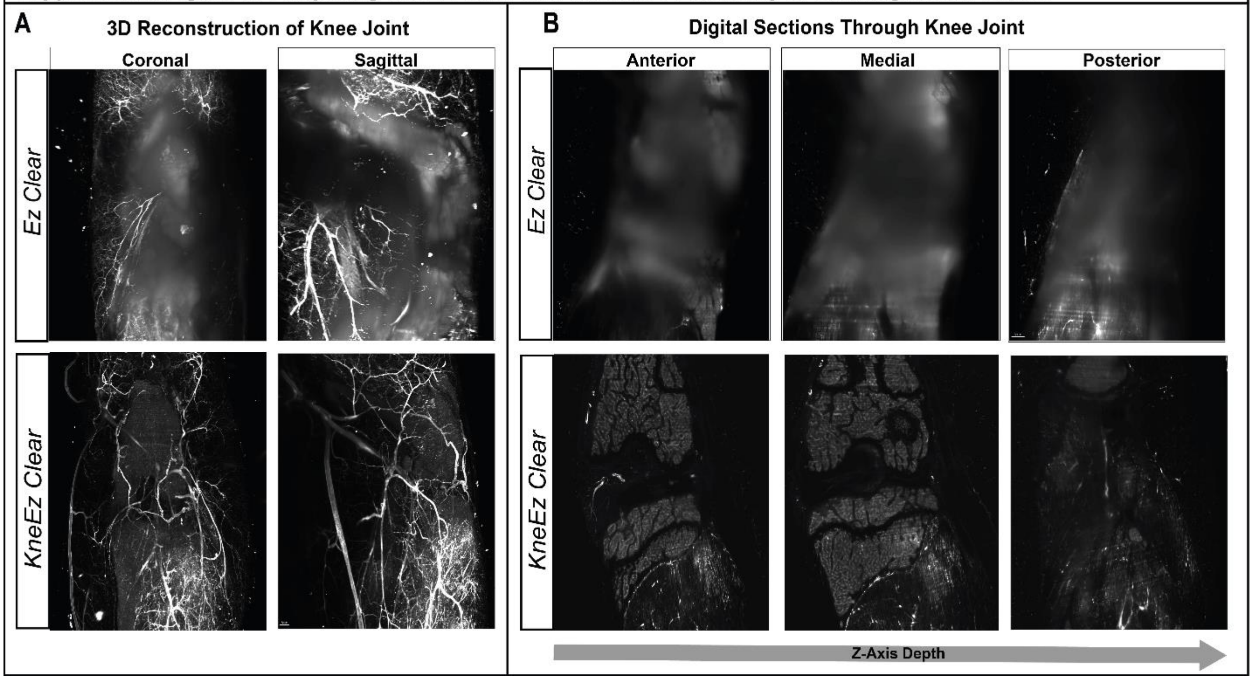


Supplemental Figure 2. Vascular Signal is Increased with a Post-perfusion Lectin Pulse

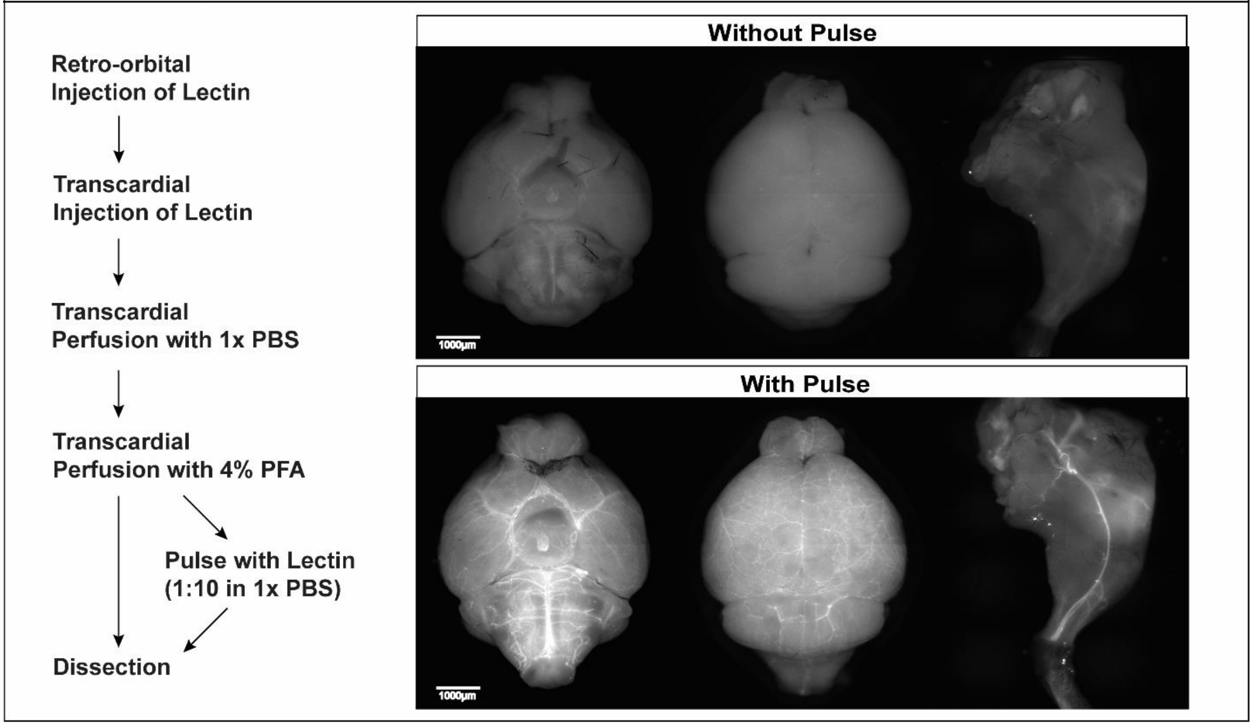

### Supplemental Figure 3. Parameters Tested for Clearing Optimization

**A**

| Fixation Temperature | Fixation Duration | Decalcification Concentration | Decalcification Duration | Heme Removal | Lipid Removal |
|---|---|---|---|---|---|
| 4°C | 16 Hours | 10% | 2 Days | $H_2O_2$ | Single Incubation |
| 21°C | 24 Hours | 14% | 1 Week | Amino Alcohol | Multistep Gradient |
| | 48 Hours | | 2 Weeks | | |

**B**

Parameters that Decreased Clearing Efficiency

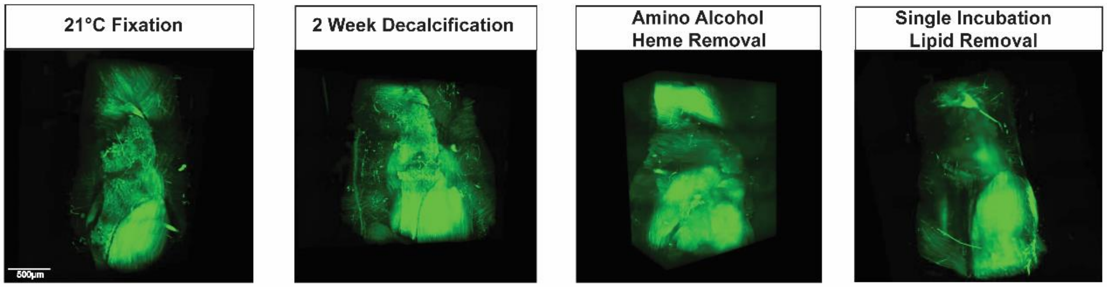

### Supplemental Figure 4. Comparison of EZ Clear and KneEZ Clear Protocols

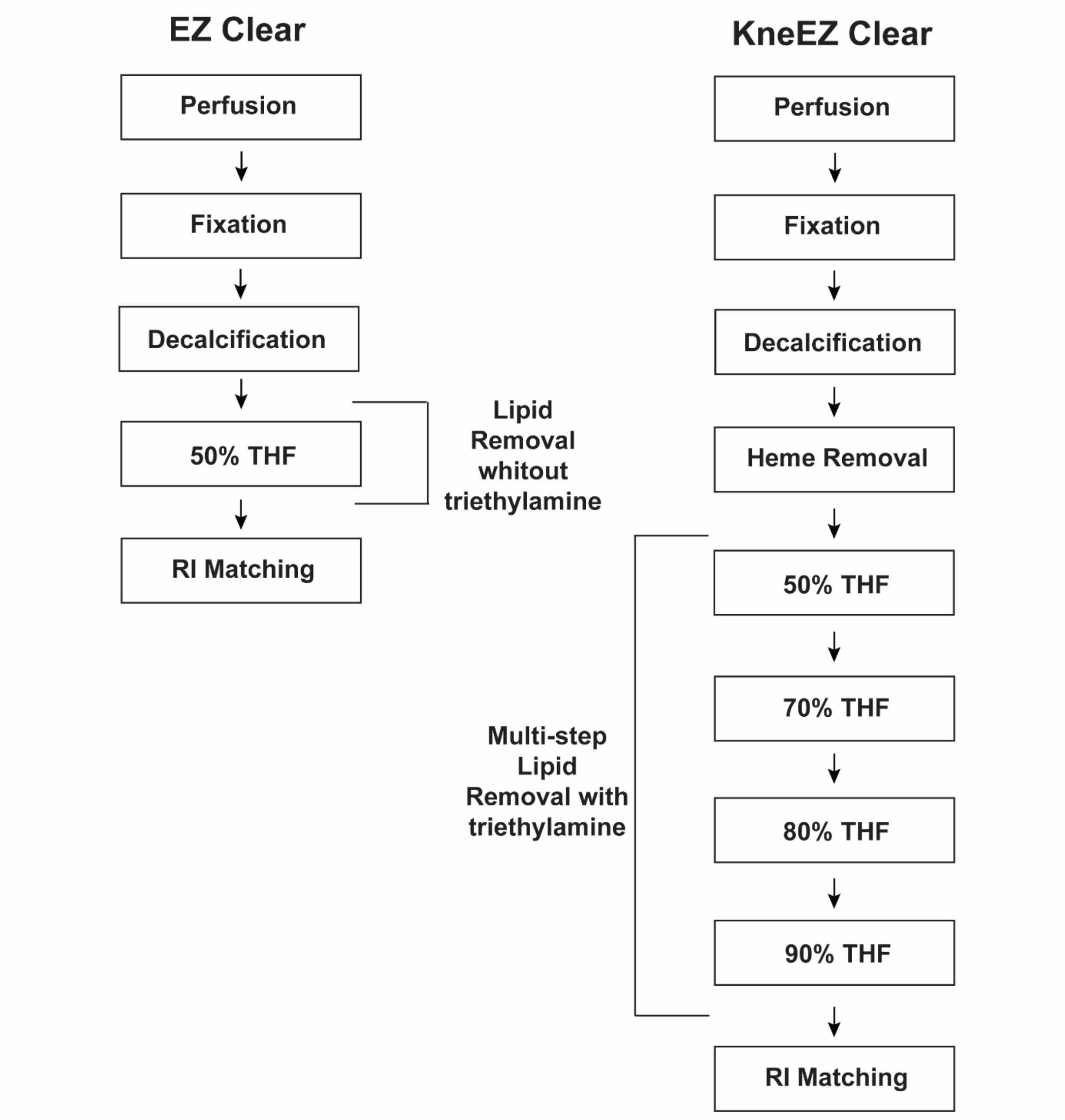

Supplemental Figure 5. Post-processing of KneEZ Cleared Knee Joints Using Cryosection-based Histology

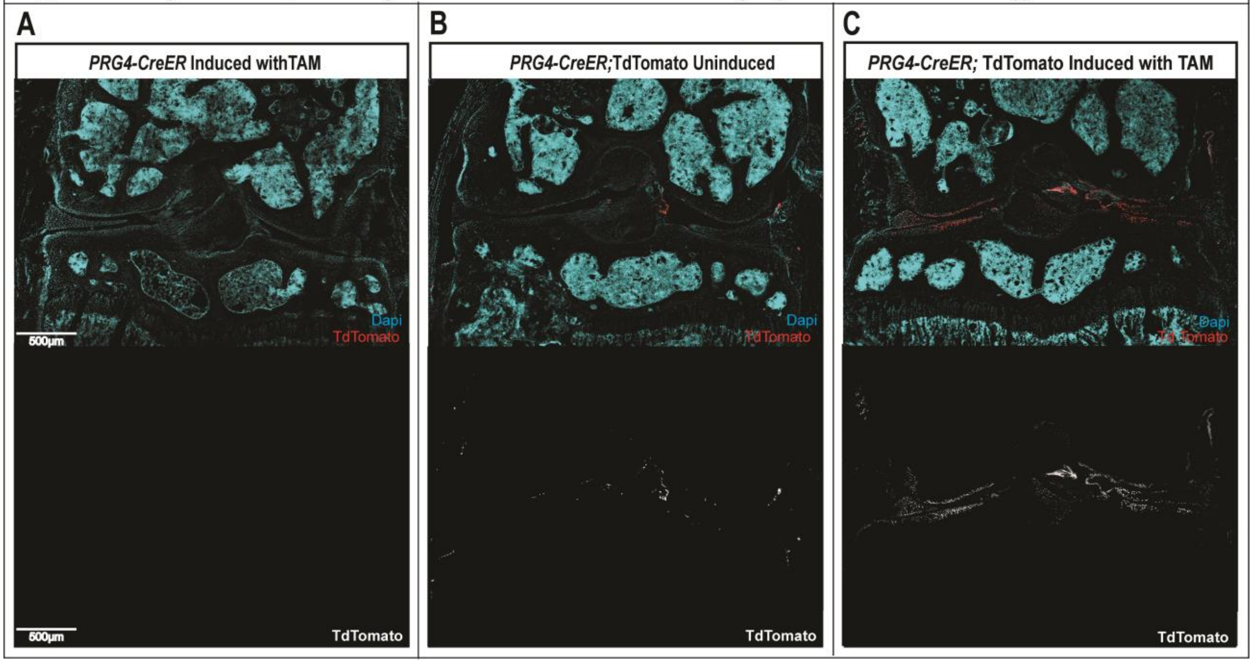

Supplemental Figure 6. Immunolabeling of Sectioned KneEZ Cleared Knee joints

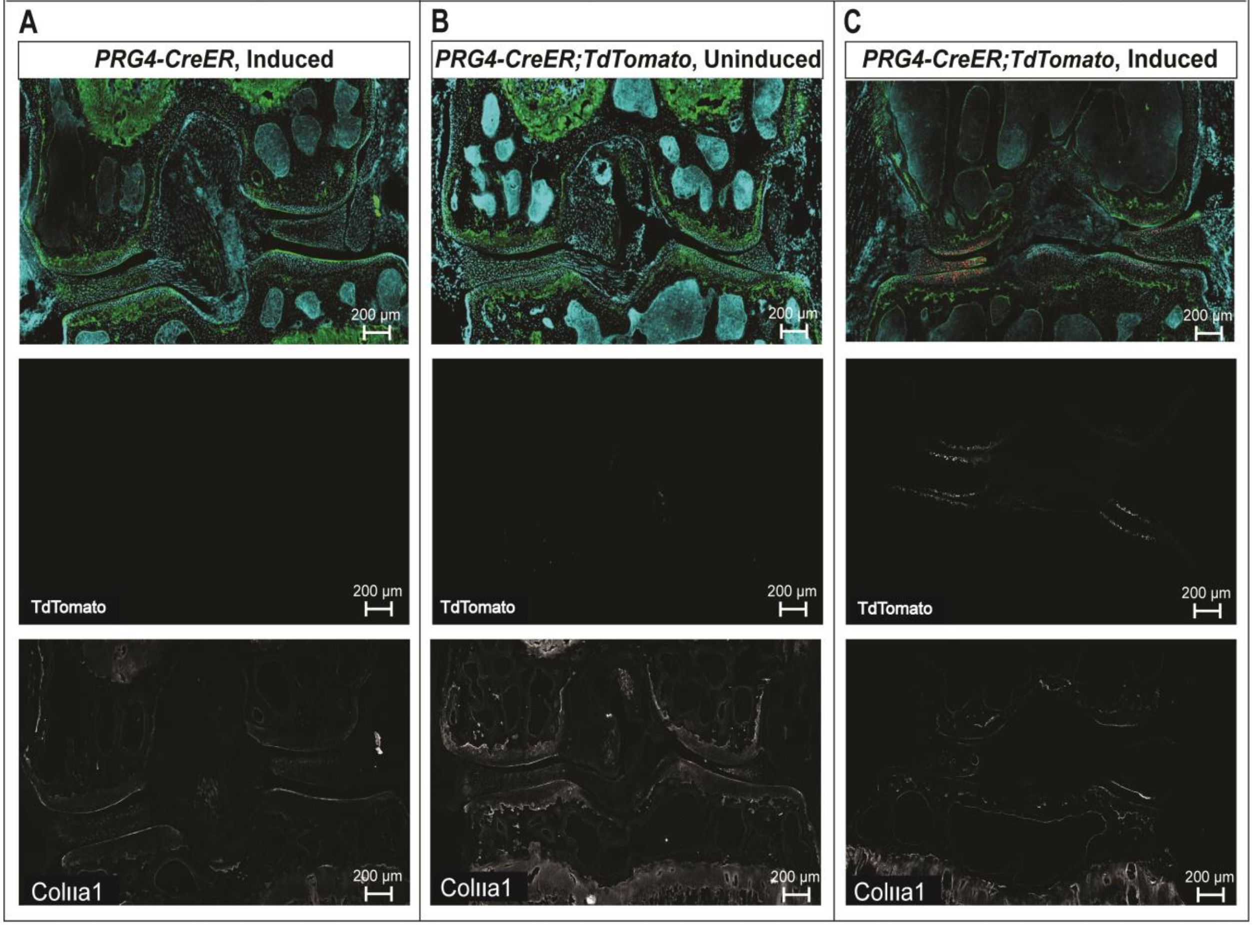


Supplemental Figure 7. Reflective Light Reveals Bone and Soft Tissue Phenotype in *Scx-Cre, FKBP10* Mice

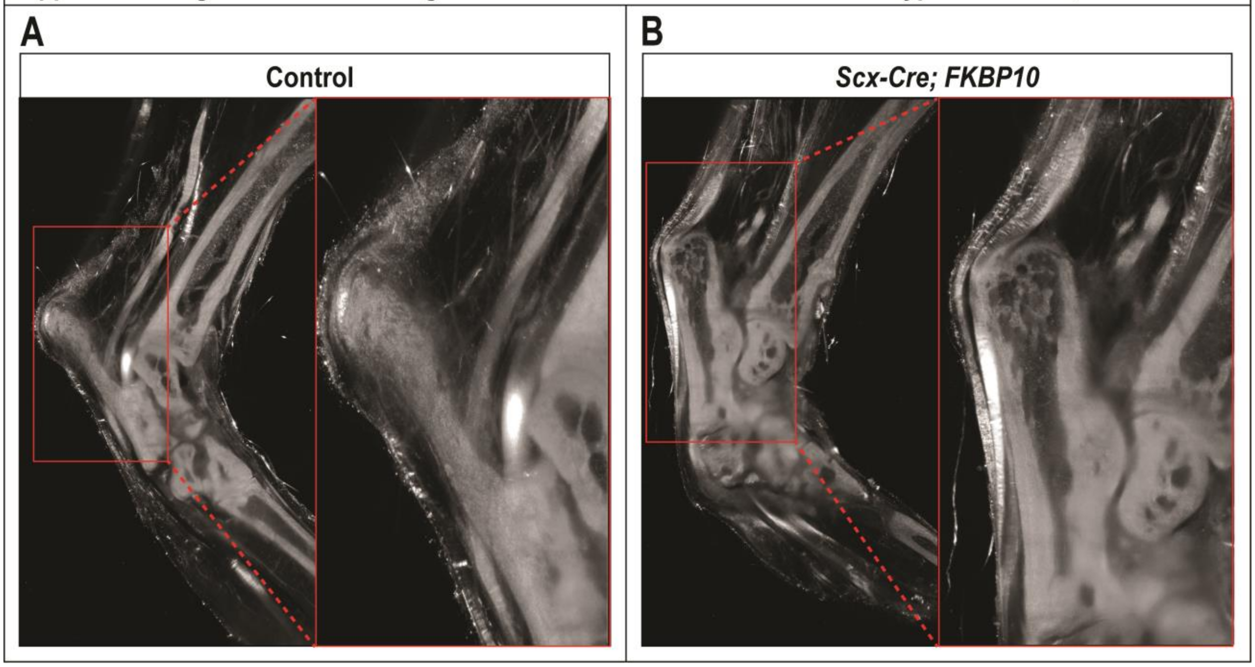
